# Chigno/CG11180 and SUMO are Chinmo-Interacting Proteins with a Role in *Drosophila* Testes Stem Cells

**DOI:** 10.1101/2021.11.03.467147

**Authors:** Leanna Rinehart, Wendy E. Stewart, Natalie Luffman, Matthew Wawersik, Oliver Kerscher

**Author notes:** Joint corresponding authors: Matthew Wawersik and Oliver Kerscher, email address.

## Abstract

Maintenance of sexual identity on the cellular level ensures the proper function of sexually dimorphic genes expressed in the brain and gonads. Disruption of genes that regulate sex maintenance alters the cellular structure of these tissues and leads to infertility and diseases, such as diabetes, obesity, and gonadal cancers. Sex maintenance in the testis of *Drosophila melanogaster* depends on the previously identified gene *chinmo* (Chronologically inappropriate morphogenesis). Chinmo’s effect on testis differentiation has been investigated in detail, but there is still much to be elucidated about its structure, function, and interactions with other proteins. Using a two-hybrid screen, we find that Chinmo interacts with itself, the small ubiquitin-like modifier SUMO, the novel protein CG11180, and four other proteins (CG4318, Ova (Ovaries absent), Taf3 (TBP-associated factor 3), and CG18269). Since both Chinmo and CG11180 contain sumoylation sites and SUMO-interacting motifs (SIMs), we analyzed their interaction in more detail. Using site-directed mutagenesis of a unique SIM in CG11180, we demonstrate that Chinmo’s interaction with CG11180 is SUMO-dependent. Furthermore, to assess the functional relevance of both SUMO and CG11180, we performed RNAi-mediated knockdown of both proteins in somatic cells of the *Drosophila* testis. Using this approach, we find that CG11180 and SUMO are required in somatic cells of adult testes, and that reduction of either protein causes formation of germ cell tumors. Overall, our work indicates that SUMO functionally links Chinmo and CG11180 in somatic cells of the adult *Drosophila* testis. Consistent with the CG11180 knockdown phenotype in male testes, and to underscore its connection to Chinmo, we propose the name Childless Gambino (Chigno) for CG11180.

## INTRODUCTION

Sex determination, or the assignment of sex, is vital for fertility and the proper function of tissues showing sexual dimorphism, including the brain and gonads (Grmai et al., 2018; Hudry et al., 2016; Mauvais-Jarvis, 2015; Sacher et al., 2013; X. Yang et al., 2006). Beyond determination of sexual identity, many animals must also maintain sexspecific transcriptional programs throughout their lifetime to ensure continued tissue homeostasis (Grmai et al., 2018; Ma et al., 2014; Matson et al., 2011; Mauvais-Jarvis, 2015; Uhlenhaut et al., 2009). *Drosophila* testes are an excellent model for understanding sex-specific stem cell regulation. Each testis is a coiled tube that includes a stem cell niche made up of hub cells, germline stem cells (GSCs), and somatic cyst stem cells (CySCs). The hub is a cluster of quiescent somatic cells at the testis apex that GSCs attach to. CySCs flank individual GSCs and provide cues for GSC self-renewal alongside hub signaling. As part of this renewal, both GSCs and CySCs divide asymmetrically to produce two daughter cells; one of which remains at the niche and retains stem cell identity while the other is displaced from the niche and initiates differentiation into either a sperm or a spermatogenesis supporting somatic cyst cell (Matunis et al., 2012).

Although upstream regulators of sex determination in *Drosophila* differ substantially from the hormonal cues that determine sex in mammals, downstream control converges on the *DSX/mab-3 related transcription factor* (Dmrt) (Clough et al., 2014; Lindeman et al., 2015). Therefore, studying *dsx* and its upstream regulators in *Drosophila* provides valuable information about sex determination and maintenance that also applies to other organisms. One upstream regulator of *dsx* within the *Drosophila* sex maintenance cascade is the protein Chinmo (*Ch*ronologically *in*appropriate *mo*rphogenesis). Chinmo, which is conserved as ZFP509/ZNF509 in mammals, is expressed in male but not female gonadal tissues, and functions downstream of the Janus kinase-Signal Transducer and Activator of Transcription (Jak/STAT) pathway required for testis stem cell self-renewal and the regulation of male germline sexual development (Flaherty et al., 2010; Kiger et al., 2001; Tulina & Matunis, 2001; Wawersik et al., 2005). Chinmo also plays a vital role in the determination and maintenance of male sexual identity in CySCs. When Chinmo expression is reduced in somatic cells of the adult testis, somatic CySCs undergo a male-to-female sex transformation, resulting in infertility and germ cell tumors (Grmai et al., 2018; Ma et al., 2014). Therefore, studying Chinmo’s structure and function is crucial to furthering our understanding of mechanisms controlling sex-maintenance in *Drosophila*, the roles of ZFP509 and other putative Chinmo orthologs in mammals, as well as function of conserved DMRT proteins (Flaherty et al., 2010). Such studies may also help us understand mechanisms by which Chinmo controls other processes such as the prevention of tumor formation, and regulation of neuronal temporal identity and eye development (Flaherty et al., 2010; Kao et al., 2012; Zhu et al., 2006).

Inspection of the Chinmo cDNA and ORF reveals several notable features. Genomic analyses indicate the existence of six distinct *chinmo* transcripts encoding a 604 amino acid and an 840 amino acid protein, respectively (flybase.org). Chinmo is a member of the BTB-ZF (broad-complex, tramtrack and bric-à-brac - zinc finger) protein family, which frequently plays a role in transcriptional regulation (Siggs & Beutler, 2012). In mammalian BTB-ZF proteins, the ZF domain determines sequence specificity while the BTB domain promotes oligomerization and the recruitment of transcriptional repressors (Siggs & Beutler, 2012). The various mammalian BTB-ZF domain proteins are involved in lymphocyte development, fertility regulation, skeletal morphogenesis, and neural development. In *Drosophila*, BTB-ZF genes other than *chinmo*, such as *fruitless, abrupt*, and *lola*, are involved in neurological development and control a variety of processes, such as cell differentiation, organ formation, and sex-specific behavior (Ito et al., 1996; Sato et al., 2019; Zhu et al., 2006). Chinmo also contains several putative motifs for covalent or noncovalent interaction with the post-translational modifier SUMO (Small Ubiquitin-like Modifier) (Kerscher, 2007). SUMO can be covalently linked to specific target proteins to modify their activity, localization, interactions, and half-life (Kerscher, 2007; Kerscher et al., 2006). Functionally, proteins that undergo modification by SUMO perform essential roles including those in transcription, mRNA processing, DNA replication, intracellular transport, and DNA-damage response (Hendriks & Vertegaal, 2016; Hochstrasser, 2009; Impens et al., 2014). The sumoylation sites found in Chinmo thus raise the possibility that SUMO targets Chinmo for modification and regulates its interactions and essential functions.

Here, we show that Chinmo interacts with itself, SUMO, and five other proteins, 3 of which are novel in function (CG4318, Ova (Ovaries absent), Taf3 (TBP-associated factor 3), CG18269, and CG11180). CG11180 encodes an uncharacterized *Drosophila* protein orthologous to human PINX1, a protein playing a vital role in the maintenance of telomere length and chromosome stability. CG11180 was identified six times in our Chinmo interactor screen and thus was chosen for further analysis. We show that the interaction between Chinmo and CG11180 depends on SUMO. Furthermore, Chinmo’s interaction with both CG11180 and SUMO are functionally relevant *in vivo* as knockdown of either of these two proteins in somatic cells of the adult *Drosophila* testis yields germ cell tumor phenotypes similar to those observed after somatic *chinmo* knockdown. Due to its interactions with Chinmo and its tumor associated germ cell differentiation defects, we propose to name CG11180, *Childless Gambino (Chigno)*. Together, our data suggest that SUMO may functionally link the activity of Chinmo and CG11180/Chigno in somatic cells of the adult *Drosophila* testis, thereby playing a key role in male fertility and spermatogenic differentiation.

## MATERIALS AND METHODS

### Yeast two-hybrid assays

Two-hybrid screening and assays were performed as described in the Matchmaker^™^ Pretransformed Libraries User Manual (Takara PT3183-1) using a Drosophila Mate & Plate^™^ Library - Universal Drosophila (Normalized A+ RNA isolated from embryo (~20 hr), larval, and adult stage *Drosophila melanogaster*) Cat. No. 630485. Briefly, a fusion of Chinmo^(native)^ with the Gal4 DNA Binding domain (DB) was expressed in yeast (AH109) and then mated to yeast two-hybrid library of normalized *Drosophila melanogaster* cDNA clones transformed in a MATa GAL4 reporter strain (Y187). Chinmo interactors were identified as colonies that robustly activated the *ADE2*, *HIS3* reporter genes and formed blue colonies on SD/-Ade/-His/-Leu/-Trp/X-alpha-Gal media. Individual clones were isolated, sequenced, and retested (see Table 1C).

### Protein domain prediction

Prediction of SIMs and sumoylation consensus sites was done using GPS SUMO (http://sumosp.biocuckoo.org/online.php). Only high confidence SIMs and sumoylation sites are reported. (Zhao et al., 2014). Other domains in Chinmo and CG1180/Chigno were mapped using https://prosite.expasy.org (Sigrist et al., 2013). Sequences and transcript analyses were retrieved from flybase.org (Larkin et al., 2021).

### Cloning and site-directed mutagenesis

BD-Chinmo clones were derived using a commercially available cDNA clone (SD04616) as template and PCR primers specific for the long and short Chinmo open-reading frame. The resulting Chinmo amplicons were recombined into two-hybrid plasmid pOBD2 as described (http://depts.washington.edu/sfields/protocols/pOBD2.html). Mutagenic primers and the Q5 site-directed mutagenesis kit (NEB E0554S) were used to generate the BD-Chinmo^non-stop^ and the CG11180/Chigno-sim* mutant. All clones were confirmed using sequence analysis and, in the case of Chinmo, western blotting (data not shown).

### Fly stocks

*c587-Gal4* (Kai & Spradling, 2003) and *eyaA3-Gal4* (Leatherman & Dinardo, 2008) were used to drive UAS-transgene expression in the somatic CySC lineage. UAS-lines include: UAS-CG11180-RNAi (Bloomington stock #28629), UAS-Smt3-RNAi (Bloomington stock #28034), UAS-CG5694-RNAi (Bloomington stock #62485), UAS-4318-RNAi (Bloomington stock #51720), UAS-Bip2/Taf3-RNAi (Bloomington stock #43174). *y,w^1118^* flies were used as wild type controls. Fly stocks were obtained from Bloomington Stock Center (http://flystocks.bio.indiana.edu/) unless otherwise specified.

### Testis collection & immunostaining

Flies were raised at 25°C unless otherwise indicated. To prevent Gal4-UAS activity during development, flies were reared at 18°C and allowed to eclose as adults for 48-72 hours, after which flies were shifted to 25°C. Adult testes were dissected, fixed and immunostained at designated ages as described (Matunis et al., 1997). Antibodies used for immunostaining were Rabbit anti-Vasa at 1:150 (R. Lehmann) or Rat anti-Vasa at 1:60 (A. Spradling/Williams, D.; DSHB), and goat anti-rabbit Alexa 568 (Invitrogen; A-11011) or goat anti-rat RRX (Jackson Labs; 112-295-068) at 1:500. Nuclei were stained using DAPI at 1 μg/mL (Roche) for 3 min.

### Microscopy & phenotype classification

Samples were mounted in 70% glycerol containing 2.5% DABCO (Sigma) and *p*-phenylenediamine anti-fade agent (Sigma) at a final concentration of 0.2 mg/mL. Slides were viewed with an Olumpus BX51 microscope equipped with a DSU spinning disc confocal system and Q-imaging RETIGA-SRV CCD camera. Images were captured and analyzed with Slidebook software by 3I. For scoring of phenotype severity, wild type testes with high-level DAPI staining restricted to the testis apex where less differentiated germ cells reside (Tulina & Matunis, 2001) were scored as normal. An increase in DAPI-bright region extending up to 10% the length of the testes coil was scored as mild, 10-30% scored as moderate, and greater than 30% of the testes coil scored as severe.

Testes with expansion of less differentiated DAPI-bright germ cells but lacking an obvious testes coil were scored as severe aberrant/underdeveloped.

## RESULTS

### Identification of Chinmo interactors

Chinmo contains a BTB protein interaction domain (amino acids (aa) 32-98), two Zinc Finger Domains (aa 517-545 and 545-573), three putative SUMO-interacting motifs (SIM) (aa 71-75, 773-777, 780-784), and at least 5 SUMO consensus sites (K288, K414, K447, K476, K779) (Fig. 1A). These domains and motifs raise the possibility that Chinmo interacts with multiple different proteins, (e.g., other Chinmo proteins or transcriptional repressors), SUMO, and sumoylated proteins. However, additional analyses are required to identify Chinmo interacting proteins that could provide functional clues. Therefore, we used a two-hybrid screen to identify these proteins and investigate Chinmo’s interactions in detail. To perform this analysis, we obtained a Chinmo 4259 nucleotide (nt) cDNA (SD04616) to clone the Chinmo ORF in frame with the Gal4 DNA Binding domain (BD) into the pOBD2 two-hybrid vector. Inspection of the Chinmo coding sequence in this cDNA clone revealed an 1815 nt ORF, terminating in a single STOP codon. This ORF translates into a 604 aa Chinmo protein containing all indicated protein interaction domains and motifs (Fig. 1A). Curiously, we noticed that omission of the STOP codon (potentially by read-through) yields a 2529 nt ORF encoding a previously reported 842 aa long Chinmo protein isoform (http://flybase.org/reports/FBgn0034528). Hence, we generated three BD-Chinmo clones: BD-Chinmo^native^ (with the Stop at nt 1813), BD-Chinmo^non-stop^ (lacking the Stop codon at position 1813), and BD-Chinmo^STOP^ (omitting sequences after the Stop codon at position 1813) (Fig. S1A). Western blots showed that all versions of BD-Chinmo expressed fusion proteins were of the expected size (data not shown).

**Fig. 1:**
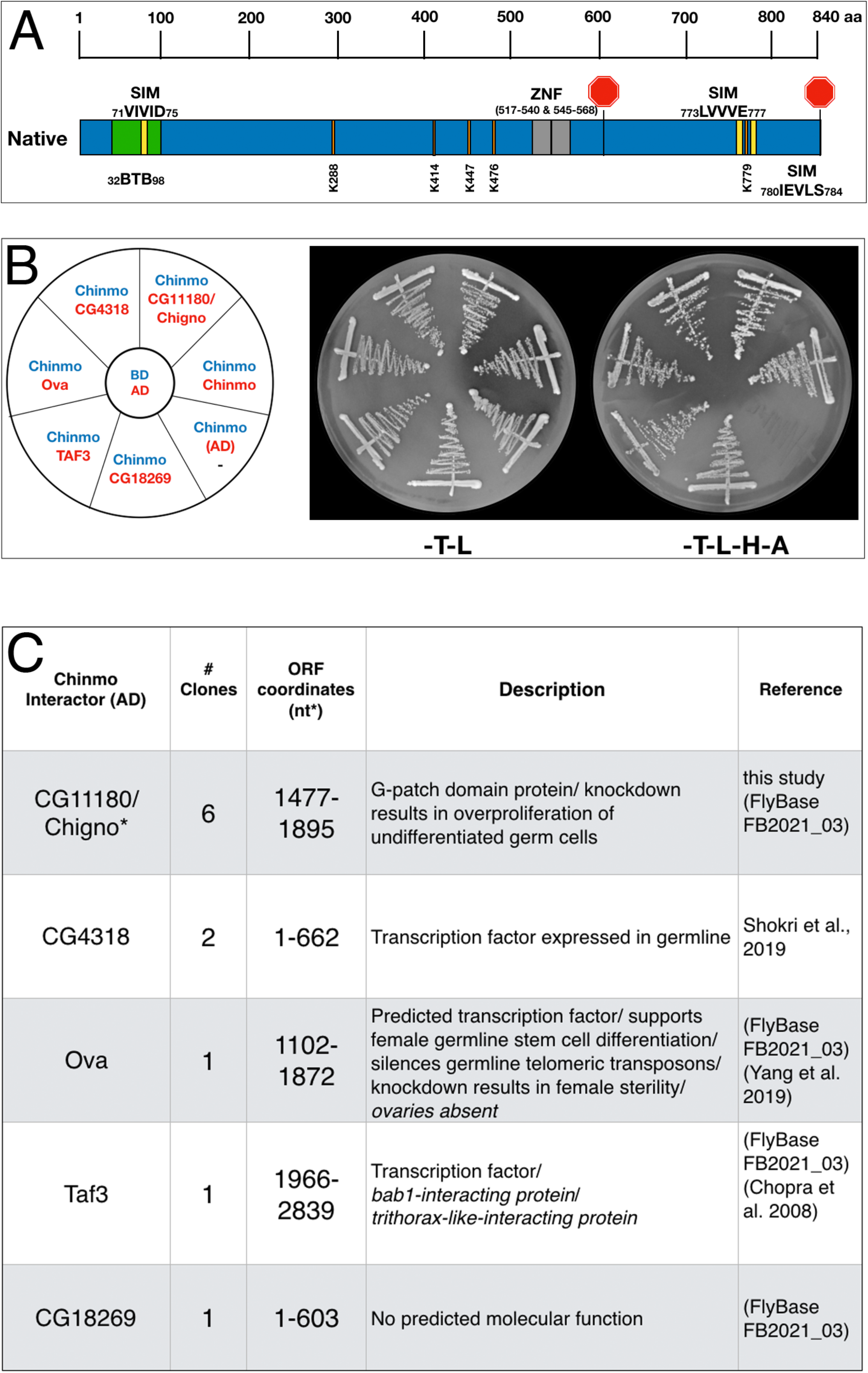
Chinmo protein domains and interactors: **(A)** Chinmo contains a BTB protein interaction domain spanning residues 32 to 98 and two zinc finger domains (ZNF) spanning from 517 to 540 and 545 to 568. Two stop codons in the cDNA terminate translation at residue 604, or, if read-through occurs, at residue 840. Furthermore, SUMO interaction motifs and sumoylation sites are shown in yellow and orange respectively. Note that a putative SIM domain in Chinmo is contained within the BTB protein-interaction domain. **(B)** Two-hybrid analysis shows that Chinmo interacts with five novel proteins (encoded by CG4318, Ova (Ovaries absent), Taf3 (TBP-associated factor 3), CG11180, and CG18269) and other Chinmo proteins. <u>Left:</u> graphic representation of BD-Chinmo (blue) and its interactors (red). BD-Chinmo/AD-empty vector is a negative control. <u>Middle</u>: The presence of BD-Chinmo (in pOBD2/TRP1) and the indicated AD constructs (in pOAD/LEU2) was confirmed by growth on growth media lacking tryptophan and leucine (-T-L). <u>Right</u>: The interaction between Chinmo constructs and Chinmo interactors is confirmed by growth on media lacking lacking tryptophan, leucine, histidine, and adenine (T-L-H-A). Plates were imaged after three days of growth. **(C)** Shown are the gene/protein name, approximate coordinates of the cDNA fragment identified, description, and reference.

To identify Chinmo interacting proteins, we used the BD-Chinmo^native^ clone to screen a two-hybrid library of *D. melanogaster* cDNAs fused to the Gal4 Activation Domain (AD-cDNA) (See materials and methods). In this screen, we identified 5 AD-cDNA fusions (CG11180, CG4318, Ova, Taf3, and CG18269). These constructs resulted in reporter gene activation (*ADE2*, *HIS3*) when co-expressed with BD-Chinmo and robust growth on media lacking adenine and histidine (Fig. 1B). As a negative control, BD-Chinmo alone (Chinmo/AD) did not result in reporter gene activation as expected (see Fig. 1B and C). Two Chinmo-interactors identified in our screen (CG4318 and Ova) are transcription factors functionally linked to germline differentiation (Yang et al., 2019); (FlyBase.org), and Taf3 is involved in transcriptional activation and regulation (Chopra et al., 2008). CG11180 and CG18269 encode proteins of unknown function (Fig.1C). While an interaction between CG4318 and Chinmo was previously identified in a high throughput interactome study (Shokri et al., 2019), interactions between Chinmo and the other four proteins are novel.

As part of our screen for Chinmo interactors, we also included an AD-Chinmo clone and found that it interacts with the BD-Chinmo^native^ clone used for library screening. In support of choosing BD-Chinmo^native^ for our screen, we did not discern a difference in the interaction between the different BD-Chinmo clones (Native, Non-Stop, and Stop) and AD-Chinmo (Fig. S1B). In summary, our 2-hybrid screen identified 5 novel Chinmo-interacting proteins and also revealed that Chinmo dimerizes with itself.

### Chinmo interaction with SUMO

Because Chinmo has several SIMs (Figure 1A) and we did not identify SUMO in our screen, we also directly assayed Chinmo’s predicted interaction with SUMO.

SUMO/Smt3 is a structurally highly conserved protein with the yeast and *Drosophila* orthologs bearing 42% sequence identity and 57% similarity. Thus, we tested the interaction of ScSmt3 from yeast (AD-SUMO) with all three BD-Chinmo variants. In our assay, AD-SUMO interacted with all three BD-Chinmo variants (BD-Chinmo^native^, BD-Chinmo^non-stop^, and BD-Chinmo^STOP^) on media lacking histidine. BD-Slx5, a SUMO interacting protein with multiple SIMs, served as a positive control and as expected, AD-SUMO alone (empty/SUMO) did not result in reporter gene activation. Using a more stringent selection (SD-His-Ade) recapitulated our finding that SUMO interacts robustly with BD-Chinmo^native^ and BD-Chinmo^STOP^. However, the interaction of Chinmo^non-stop^ with SUMO was weaker, as apparent from reduced growth on SD-His-Ade media (Fig. 2A). An ONPG ß-galactosidase assay to quantitate the interaction of SUMO and Chinmo also showed that the STOP and Native Chinmo clones showed equivalent reporter gene activation, while the interaction of AD-SUMO and the Chinmo^non-stop^ was reduced to background levels (Fig. 2B). In summary, our data show that both BD-Chinmo^native^ and BD-Chinmo^STOP^ interact with SUMO, and that this interaction may depend on the SIM (aa 71-75) in Chinmo.

**Fig. 2:**
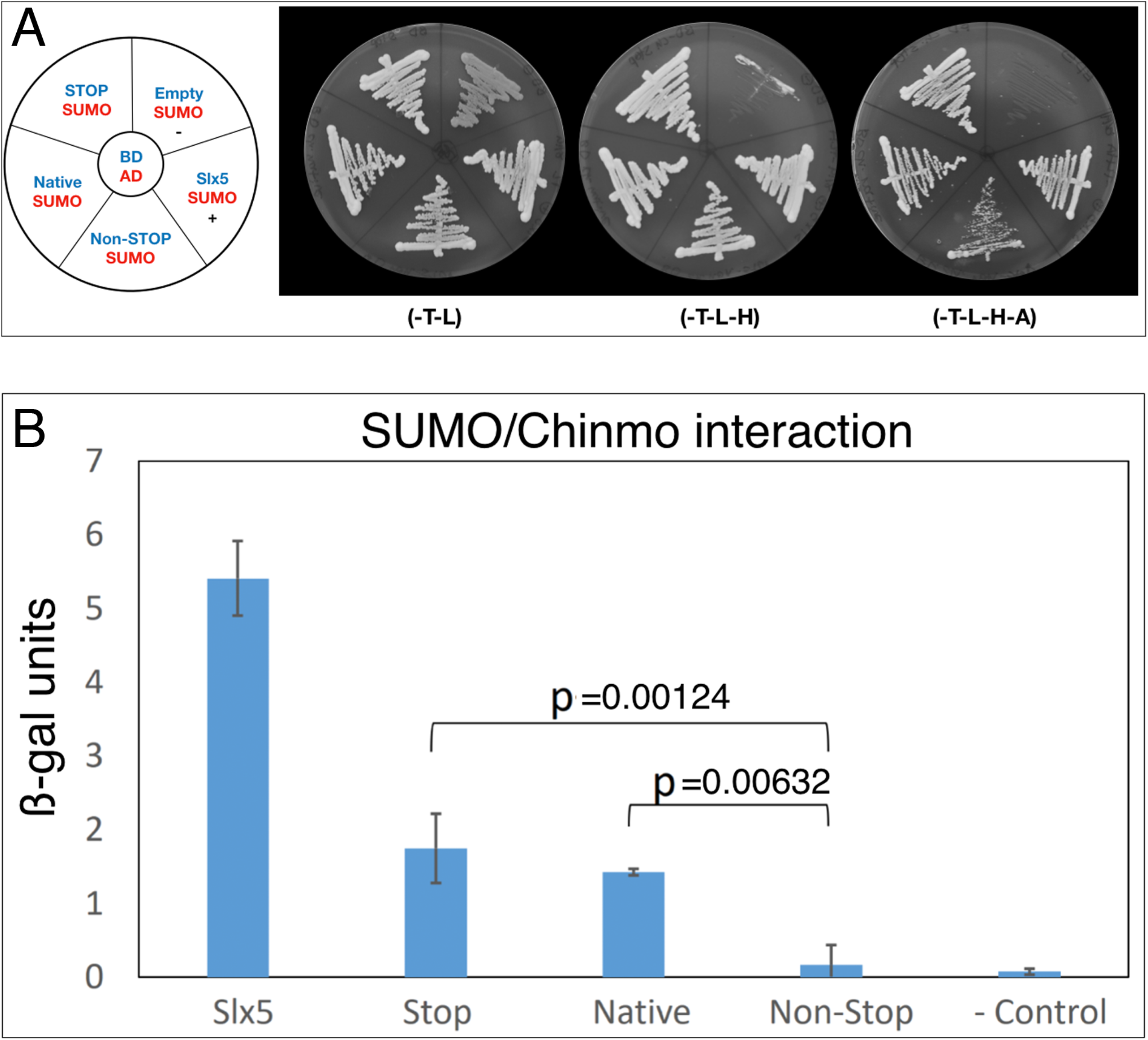
Chinmo interacts with SUMO. **(A)** Two-hybrid analysis shows that BD-Chinmo Native and BD-Chinmo Stop (and to a lesser degree BD-Chinmo Non-Stop) interact with AD-SUMO. <u>Left</u>: Graphic representation of the three BD-Chinmo clones (Native, Stop, and Non-Stop) (blue) and AD-SUMO (red). Negative (BD-empty vector with AD-SUMO) and positive controls (BD-Slx5 with AD-SUMO) are as indicated. <u>Middle</u>: The presence of the indicated BD-Chinmo constructs (in pOBD2/TRP1) and AD-SUMO (in pOAD/LEU2) was confirmed by growth on growth media lacking tryptophan and leucine (-T-L). <u>Right</u>: Interactions are confirmed by growth on media lacking tryptophan, leucine, histidine, and adenine (T-L-H and -T-L-H-A). Plates were imaged after three days of growth. **(B)** Quantification of the interaction of SUMO with the indicated Chinmo proteins using a quantitative ONPG assays and strains shown in A. Tukey Kramer multiple comparison procedure was used to determine significant differences in SUMO interaction.

### The interaction of Chinmo with CG11180/Chigno depends on SUMO

Both Chinmo and the partial cDNA of CG11180/Chigno (aa 373-621) isolated in our screen contain SIM domains (Fig 3A). Therefore, we also assessed if Chinmo’s interaction with CG11180/Chigno is SUMO-dependent. For this analysis, we focused on the functional role of a SIM mutant (sim*) in AD-CG11180/Chigno for two reasons: (i) One of Chinmo’s SIMs resides within the BTB domain and a sim* mutant may affect its dimerization, and (ii) Chinmo has multiple sumoylation sites that potentially interact with the SIM in CG11180/Chigno (unlike the Chigno clone). For our analyses, we created an AD-CG11180/Chigno^sim*^ mutant by site-directed mutagenesis, replacing the SIM in CG11180/Chigno (VIVID) with alanine’s (AAAAA). We then tested both the AD-CG11180/Chigno^sim*^ mutant and the AD-CG11180/Chigno clone for the ability to interact with BD-SUMO, BD-Chinmo, and an empty BD control plasmid. We found that AD-CG11180/Chigno interacts with both BD-SUMO and BD-Chinmo. In contrast, the AD-CG11180/Chigno^sim*^ mutant failed to interact with both BD-SUMO and BD-Chinmo (Fig. 3B). As a negative control, the empty BD vector failed to elicit reporter gene activation with both the AD-CG11180/Chigno^sim*^ mutant and the AD-CG11180/Chigno clone. Additionally, the positive control (p53/T) showed reporter gene activation, as expected. Together, these results suggest that the VIVID sequence in CG11180/Chigno constitutes a *bona-fide* SIM that is required for the interaction with Chinmo.

**Fig. 3:**
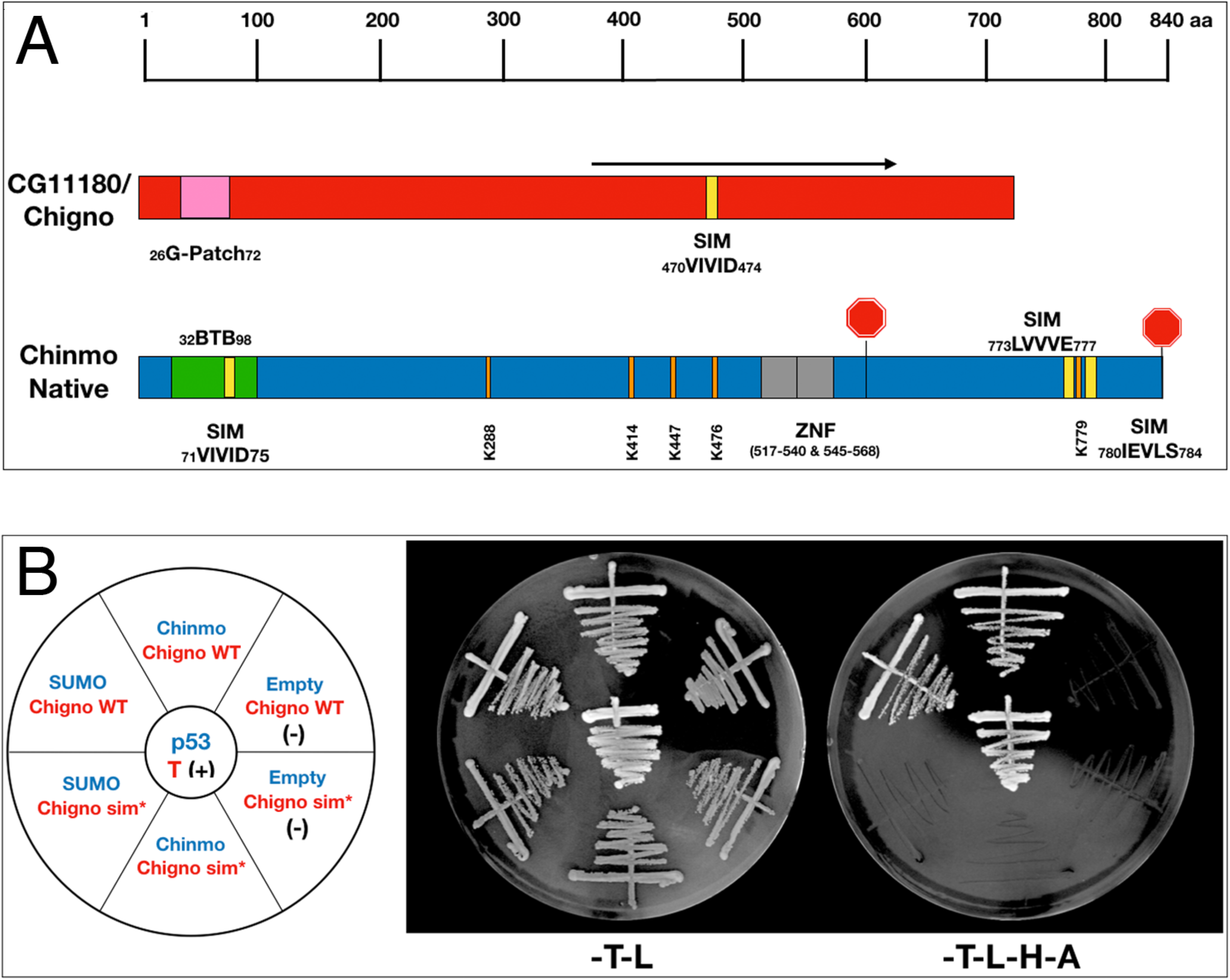
Chinmo’s interaction with CG11180/Chigno depends on a SIM in Chigno. **(A)** Comparison of protein domains and motifs in CG11180/Chigno (red) and Chinmo (blue). Coordinates of SIMs and sumoylation consensus sites as shown. Note that the SIM domain in Chinmo is contained within the BTB protein-interaction domain. The black arrow demarcates the CG11180/Chigno protein fragment encoded by the cDNA isolated in our screen (373-621). **(B)** <u>Left</u>: Graphic representation of BD-Chinmo constructs (blue) and AD-CG11180/Chigno, either containing or lacking the SIM at residues 71 to 75 (red). Negative controls (empty pOBD2/TRP1) and a positive control (p53/T) are as indicated. <u>Middle</u>: The presence of the indicated BD constructs (blue - pOBD2/TRP1) and either AD-CG11180/Chigno WT or AD-CG11180/Chigno sim* (pOAD/LEU2) was confirmed by growth on growth media lacking tryptophan and leucine (-T-L). <u>Right</u>: Interactions are confirmed by growth on media lacking tryptophan, leucine, histidine, and adenine (-T-L-H-A). Plates were imaged after three days of growth.

### CG11180/Chigno and SUMO are required in somatic cells of the adult *Drosophila* testis

We next assessed the functional significance of identified Chinmo-interacting proteins *in vivo*. Because Chinmo functions in somatic CySCs of adult *Drosophila* testes (Flaherty et al., 2010; Grmai et al., 2018; Ma et al., 2014), we performed RNAi knockdown of Chinmo-interacting proteins in the CySC-lineage using the Gal4-UAS system (Brand & Perrimon, 1993). Specifically, UAS-RNAi constructs against Chinmo-interactors identified through our screen, as well as SUMO/Smt3, were expressed in the CySC lineage using the c587-Gal4 driver (Kai & Spradling, 2003). Adult testes were then isolated and processed for fluorescence immunostaining. Because functional CySCs are critical for normal germ cell differentiation, and inhibition of Chinmo results in germ cell differentiation defects associated with over-proliferation of under-differentiated germ cells, testes were assayed for germ cell differentiation defects in our initial analysis presented here.

We find that somatic knockdown of CG4318, *ova*, or *Taf3* show no germ cell differentiation defects in 0-4 day old adult testes (Figure S2). Similar results were obtained for 7-12 day old testes, though *ova* knockdown caused lethality in aged flies (data not shown). However, RNAi knockdown of CG11180/Chigno and SUMO/Smt3 using the c587-Gal4 driver caused developmental lethality in flies reared at 25°C. For technical reasons, we were unable to test a UAS-RNAi construct for the impact of CG18269.

To assess functionality of CG11180/Chigno in adult testes, experiments were adapted to prevent developmental lethality. Specifically, flies carrying the somatic c587-Gal4 driver and a UAS-CG11180-RNAi construct were reared at lower temperature (18°C) to reduce Gal4 function. Upon emergence from their pupal cases (eclosion), adult male flies were then upshifted to 25°C to permit full Gal4 activation. Under these conditions, we find that knockdown of CG11180/Chigno in somatic cells of the adult testes results in formation of germ cells tumors marked by over-proliferation of undifferentiated germ cells that are not observed in controls (Figure 4). Severity of phenotype varies (Figure 4B-F), but at its most severe, testes lack their normal coiled structure and show an absence of spermatogonial differentiation (Figure 4E-F). Consistent with adult induction of RNAi knockdown after temperature upshift, phenotype severity increases with age (Figure 4G).

**Fig. 4:**
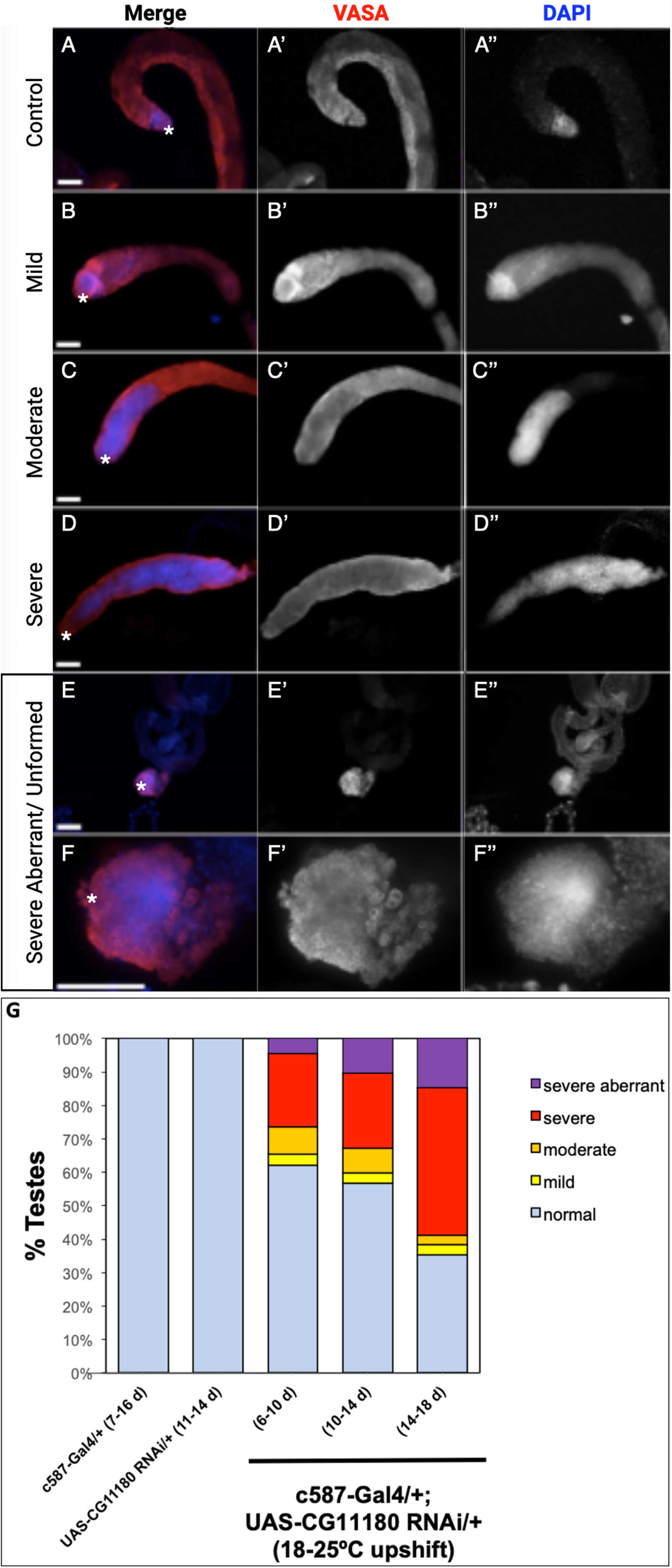
Somatic knockdown of the Chinmo interactor, CG11180/Chingo, in adult testes impacts germ cell proliferation and differentiation. **(A-F)** Adult testes 7-10 days after eclosion immunostained with anti-Vasa (red) and DAPI (blue) to reveal germ cells or nuclei respectively. Vasa and DAPI shown alone in black and white (A’-F’ and A”-F”, respectively). Testes apex containing the GSC niche indicated with an asterisk. Scale bars = 70 μm. (A) c587-Gal4/+ control testis showing presence of undifferentiated germ cells with spherical morphology and bright DAPI staining near the GSC niche and differentiating spermatogonia with dim DAPI staining extending away from the niche. (B-F) c587-Gal4/+; UAS-CG11180-RNAi/+ flies reared at 18°C then upshifted to 25°C showing defects in germ cell differentiation marked by expansion of undifferentiated germ cells that display spherical morphology and bright DAPI staining. Representative images of testes with mild (B), moderate (C), and severe (D) phenotype are shown. A severe aberrant/unformed testis at low (E) and high magnification (F) lacking an obvious testes coil while showing expansion of undifferentiated germ cells is also shown. **(G)** Graph showing percentage of control and c587-Gal4/+; UAS-CG11180-RNAi/+ testis displaying phenotype at specific ages, with phenotype severity increasing over time after somatic CG11180 knockdown (c587-Gal4/+ (7-16d), n = 56; UAS-CG11180 RNAi/+ (11-14d), n = 11; c587-Gal4/+; UAS-CG11180 RNAi/+ (6-10d), n = 77; c587-Gal4/+; UAS-CG11180 RNAi/+ (10-14d), n = 64; c587-Gal4/+; UAS-CG11180 RNAi/+ (14-18d), n = 34).

Conditions were also adapted to circumvent developmental lethality observed after SUMO/Smt3 knockdown using the c587-Gal4 driver. For these experiments, we attempted to knockdown SUMO/Smt3 using another Gal4 driver, EyaA3-Gal4 (Leatherman & Dinardo, 2008), that is also active in the somatic CySC lineage. Under these conditions, flies carrying the UAS-SUMO/Smt3-RNAi transgene were viable whether reared at 25°C, or at 18°C then upshifted to 25°C. Furthermore, under both conditions, adult males displayed a similar phenotype as CG11180/Chigno knockdown testes, with germ cell tumors marked by over-proliferation of undifferentiated germ cells (Figure 5A-B). Like CG11180/Chigno knockdown testes, this phenotype was observed at varying degrees and showed age-dependence after temperature upshift (Figure 5B-D).Taken together, these data suggest that CG11180/Chigno and SUMO/Smt3 play an important role in the regulation of somatic cell behavior in adult testes, with indirect impacts on germ cell proliferation and differentiation. Specific mechanisms by which these genes function in the testis soma, and their connection to each other as well as Chinmo, are topics for future studies.

**Fig. 5:**
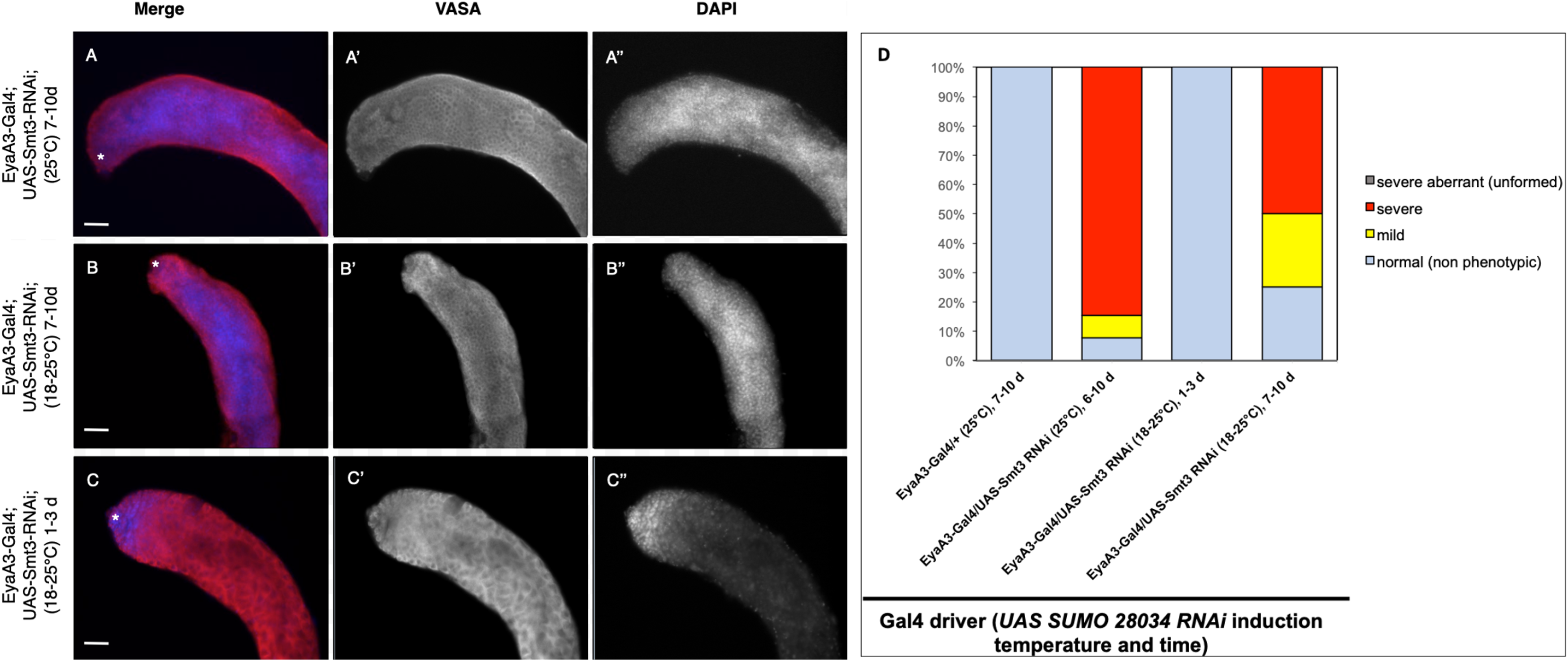
Somatic knockdown of SUMO/Smt3 in the adult testis causes proliferation of undifferentiated germ cells. **(A-C)** EyaA3-Gal4; UAS-Smt3-RNAi adult testes immunostained with anti-Vasa (red) and DAPI (blue) to detect germ cells and nuclei respectively. Vasa (A’-C’) and DAPI (A’’-C’’) are displayed individually in black and white. The testes apex containing the GSC niche is indicated with an asterisk. Scale bar = 70 μm. (A) 7-10d EyaA3-Gal4; UAS-Smt3-RNAi testis reared at 25°C displaying severe overproliferation of undifferentiated germ cells as indicated by expansion of spherical germ cells with strong DAPI fluorescence (compare to control in Fig. 4A). (B) 7-10d EyaA3-Gal4; UAS-Smt3-RNAi testis upshifted from 18 to 25°C for 7 days displaying a similar phenotype marked by overproliferation of undifferentiated germ cells. (C) 1-3d Eya A3-Gal4; UAS-Smt3-RNAi testis raised at 18°C then upshifted for 1 day is similar to controls. **(D)** Graph showing variance in phenotype severity in EyaA3-Gal4; UAS-Smt3-RNAi testes with different treatments as compared to controls (EyaA3-Gal4/+ (25°C) 7-10d, n = 23; EyaA3-Gal4/UAS-Smt3 RNAi (25°C) 6-10d, n = 13; EyaA3-Gal4/UAS-Smt RNAi (18°-25°C) 1-3d, n = 8; EyaA3-Gal4/UAS-Smt3 RNAi (18-25°C) 7-10d, n = 4).

## DISCUSSION

In this study we set out to identify Chinmo-interacting proteins using a two-hybrid screen and targeted two-hybrid assays. Our results suggest that Chinmo, a putative BTB-zinc finger transcription factor, interacts directly with three uncharacterized proteins, several transcriptional regulators, SUMO/Smt3, and itself. Chinmo dimerization is not surprising as BTB-zinc finger transcription factors are known to homo and hetero-dimerize via their BTB domains. Because most BTB-domain proteins are transcription factors, their homodimerization is predicted to increase affinity for their targets and modulate transcriptional repression properties. In contrast, some BTB domain proteins also act as substrate adapters for ubiquitin E3 ligases, targeting heterologous substrate proteins they bind for ubiquitination (Perez-Torrado et al., 2006).

Among the Chinmo-interacting proteins, CG4318, Ova, Taf3 and SUMO stand out for their previously identified roles in *Drosophila* reproductive tissues. Specifically, CG4318 encodes a putative transcription factor expressed in the germline that was previously identified as a Chinmo-interacting protein through a two-hybrid screen designed to map the *Drosophila* transcription factor interactome (Shokri et al., 2019). Identification of CG4318 in this study, therefore, validates our experimental approach. Additionally, Ova is required for fertility in female flies and functions to support ovarian germline stem cell differentiation (F. Yang et al., 2019), while Taf3 is required for the expression of germline genes in primordial germ cells (Yatsu et al., 2008) and is known to interact with proteins, like Chinmo, that contain BTB domains (Pointud et al., 2001). In the present study, we did not observe an obvious phenotype for the knockdown of CG4318, Ova, and Taf3 in somatic cells of the testis. It is, therefore, possible that these proteins interact with Chinmo in the germline, or in other tissues with Chinmo expression such as the nervous system where Chinmo regulates neuronal temporal identity during post-embryonic brain development (Yu & Lee, 2007).

Since Chinmo has multiple SUMO-interacting motifs and SUMO consensus sites, we also predicted to identify SUMO or sumoylated proteins in our screen. While we failed to isolate a *Drosophila* SUMO clone in our screen, we confirmed the interaction of Chinmo with SUMO in a two-hybrid assay with the yeast *ScSMT3*. It is, therefore, possible that the *Drosophila* cDNA prey library we used did not contain a full-length SUMO clone. Consistent with a role for *Drosophila* SUMO functioning in concert with Chinmo, SUMO and Sumoylation genes are expressed in the *Drosophila* germline during spermatogenesis (Hashiyama et al., 2009), and SUMO has been shown to play a key role in CySC regulation in *Drosophila* testes. Specifically, SUMOylation promotes CySC self-renewal via regulation of the Hedgehog (Hh) signaling pathway (Lv et al., 2016). Indeed, the Hh pathway transcription factor, Cubitus interruptus (Ci), interacts directly with the SUMO E3-ligase, Lesswright (Lwr). Additionally, somatic over-expression of mutant versions of Ci lacking SUMOylation sites results in CySC over-proliferation (Lv et al., 2016). Consistent with our observation that somatic SUMO knockdown causes over-proliferation of under-differentiated germ cells, loss of *lwr* function in somatic cells has also been shown to cause CySC over-proliferation that leads to germ cell tumor formation (Lv et al., 2016). Given that somatic Chinmo over-expression also causes CySC expansion (Flaherty et al., 2010), our observation that SUMO physically interacts with Chinmo, suggests a key role for SUMO in modulating Chinmo’s capacity to regulate CySC behavior.

Less is known about function of the novel proteins, CG18269 and CG11180/Chigno. While we did not yet assess the impact of CG18269 knockdown, there is some evidence suggesting the gene product of CG18269 may function in the testis. Specifically, CG18269 is downregulated in testis where germ cells are subjected to oxidative stress (Tan et al., 2018). Furthermore, while CG18269 expression levels are low in adult *Drosophila* tissues, RNA-Seq indicates an enrichment in the adult testis (Chintapalli et al., 2007). Thus, it is possible that CG18269 functions alongside Chinmo in the adult germline and/or somatic cells of the testis niche.

The main focus of analysis of Chinmo interactors fell on CG11180/Chigno because somatic knockdown of CG11180/Chigno caused formation of germ cell tumors. This provides evidence that CG11180/Chigno functions in CySCs where Chinmo regulates somatic sex maintenance and stem cell renewal. This notion is further supported by a whole-genome RNAi screen where CG11180/Chigno knockdown in the CySC lineage leads to male infertility marked by lack of germ cell differentiation (Fairchild et al., 2017). Further discussion of potential roles for CG11180/Chigno in adult testes and the significance of its interactions with Chinmo are presented below. A model by which SUMO may coordinate function of Chinmo-Chinmo as well Chinmo-Chigno interactions is also presented (Fig 6).

**Fig. 6:**
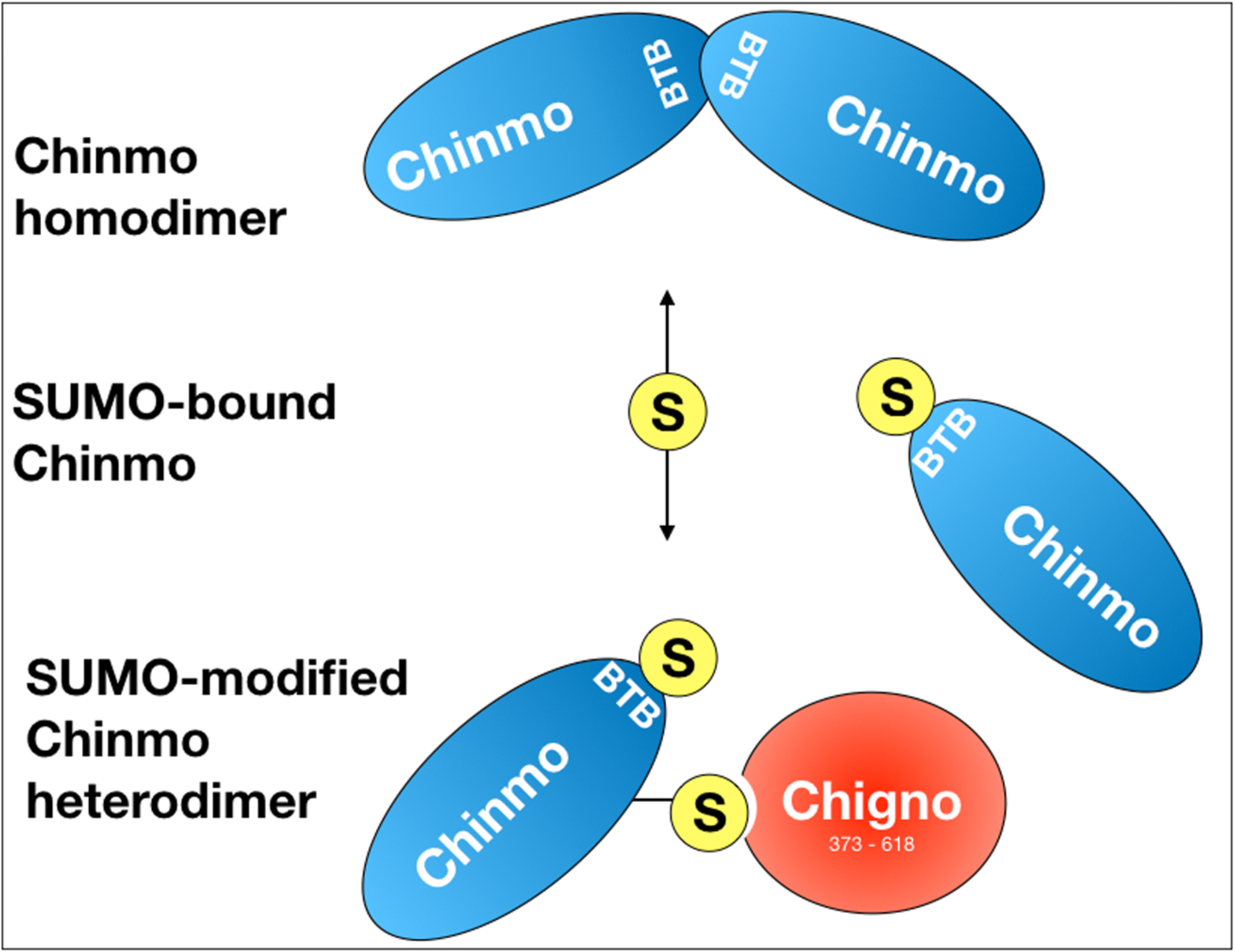
Hypothetical model of Chinmo/SUMO/Chigno interaction in testes: SUMO-binding of Chinmo (blue) prevents homodimerization and allows for interaction with Chigno and likely other SIM-containing proteins. A SIM in the BTB domain of Chinmo may prevent homodimer formation of SUMO-bound Chinmo. SUMO binding of Chinmo may then lead to conjugation of SUMO to Chinmo, involving one or several lysines.

### CG11180/Chigno function and the significance of Chinmo-Chigno interactions

As previously indicated, formation of germ cell tumors marked by germ line overproliferation and germ cell differentiation defects after somatic knockdown of CG11180/Chigno provides physiological evidence for CG11180/Chigno function in somatic cells of the testes. The direct interaction between CG11180/Chigno and Chinmo also suggests that these two proteins can function together in this tissue. Indeed, a CG11180/Chigno clone encompassing amino-acids 373-621 was identified 7 times in our 2-hybrid screen. While CG11180/Chigno is a novel protein, it contains a G-patch nucleic acid binding domain and its closest human homolog is PINX1, a potent telomerase inhibitor that interacts with TRF1, a component of the Shelterin complex (Zhou & Lu, 2001). Furthermore, PINX1 has been shown to be expressed in both somatic and germline tissues of rats, with more abundant expression in the heart, liver, and testis (Cheung et al., 2017; Oh et al., 2007). While *Drosophila* telomere replication differs dramatically from mammalian cells (Mason et al., 2008), fly telomeres are protected by a Terminin telomere-capping complex that consists of multiple proteins which share features of the Shelterin complex that interacts with PINX1 (Cicconi et al., 2017; Raffa et al., 2010). Thus, it is possible that CG11180/Chigno plays a role in telomere and/or DNA protection in somatic cells of the *Drosophila* testes. While roles for Chinmo in telomere binding have previously not been characterized, our analyses indicating direct physical interactions between CG11180/Chigno and Chinmo suggest an intriguing new role for Chinmo complexes in the regulation of cell behavior. Alternatively, as an interactome screen for proteins that modulate insulin signal suggests that CG11180/Chigno binds to the downstream transcription factors Tsc1 and Foxo (Vinayagam et al., 2016), it is also possible that CG11180/Chigno acts in concert with Chinmo to modulate the effects of insulin signaling on testes stem cell behavior (Amoyel et al., 2016).

### Role of SUMO in regulation of CG11180/Chigno and Chinmo

Sumoylation plays a pivotal role in transcriptional repression and the formation of protein complexes via protein sumoylation and SIMs (Kerscher, 2007; Wilkinson & Henley, 2010). As we have identified SUMO as a Chinmo interacting protein, it is, therefore, likely that sumoylation of Chinmo on one or several consensus site lysines regulates its activity and interactions as a transcription factor. Additionally, a bona-fide SIM is situated within the BTB dimerization domain of Chinmo. This suggests that SUMO-binding of this SIM could alter Chinmo’s conformation or interfere with its ability to dimerize and/or heterodimerize. Indeed, CG11180/Chigno also contains a SIM motif that is located at aminoacids 470-474 that facilitates binding to Chinmo; suggesting that SUMO binding might alter Chinmo-Chigno interactions. Consistent with this, we found that mutation of CG11180/Chigno’s SIM completely abolished the interaction of Chinmo with our CG11180/Chigno clone. This indicates that sumoylation of Chinmo is, in fact, required for its interaction with CG11180/Chigno. SUMO could, therefore, be utilized to recruit the Chinmo monomer to chromatin-associated CG11180/Chigno, or *vice versa*. A hypothesized model depicting the capacity of SUMO binding to modulate Chinmo-Chigno as well as Chinmo-Chinmo interactions is shown in Figure 6. Understanding whether SUMO binding mediates formation of Chinmo homodimers and/or Chinmo-Chigno interactions *in vivo*, as well as the functional significance of these interactions, remains to be determined.

## CONCLUSIONS

We hypothesized that a structure/function analysis of Chinmo and its interacting proteins will further our understanding of mechanisms controlling stem cell renewal and sex-maintenance in *Drosophila*. In summary, our study provides evidence for a complex relationship of Chinmo with other transcription factors and several proteins, such as SUMO, involved in somatic and germline regulation. Chinmo’s role in sex maintenance is restricted to testis CySCs, and our structure-function analysis combined with RNAi assays suggest that a Chinmo/Chigno/SUMO complex plays a role in fertility and prevention of testicular tumors. However, it is likely that some of the Chinmo interactions we have identified are not specific to the testis soma, but rather play important regulatory roles in germ cells or other tissues (e.g. neurons). Future studies will investigate details of the subcellular localization of Chigno, provide a more detailed structure-function analysis of the Chinmo/Chigno/SUMO complex, and focus on its precise role in the *Drosophila* testes and other tissues.

## Supporting information

Strain Table 1

Fig.S1 Rinehart

Fig.S2 Rinehart

## ACKNOWLEDGEMENTS

We are grateful to all our colleagues who have supported our labs with antibodies, stocks and technical assistance. We would also like to acknowledge the Bloomington *Drosophila* Stock Center at Indiana University for maintaining and providing fly stocks, and the Developmental Studies Hybridoma Bank developed under the auspices of the NICHD and maintained by The University of Iowa, Department of Biology. We would specifically like to thank members of the Kerscher and Wawersik labs, as well as Lidia Epp for help with sequencing.

## Supplementary Figures

**Fig. S1: Analysis of the interaction of Chinmo variants (A)** Native, Stop, and Non-Stop based on cDNA clone SD04616. Chinmo Native and Stop constructs are predicted to encode a 604 aa proteins. Chinmo Non-Stop is predicted to encode a 842 aa Chinmo protein. Domains in all contractors as indicated: BTB protein interaction domain (residue 32 to 98) and two zinc finger domains (ZNF) spanning (residues 517 to 540 and 545 to 568), as indicated. SUMO interacting motifs (SIMs) and SUMO consensus sites are shown in yellow and orange respectively. Note that the SIM domain in Chinmo is contained within the BTB protein-interaction domain. **(B)** <u>Left</u>: Graphic representation of AD and BD pairings of the three different Chinmo clones (Native, Stop, and Non-Stop). Negative (BD-empty vector/AD-Chinmo Stop) and positive controls (p53/T) are as indicated. <u>Middle</u>: The presence of the indicated BD constructs (in pOBD2/TRP1) and AD constructs (in pOAD/LEU2) was confirmed by growth on growth media lacking tryptophan and leucine (-T-L). <u>Right</u>: The interaction of the Chinmo Native, Stop, and Non-Stop with other Chinmo proteins is confirmed by growth on media lacking tryptophan, leucine, histidine, and adenine (-T-L-H-A). Plates were imaged after three days of growth.

**Fig. S2: Somatic knockdown of the Chinmo interactors, CG4318, Taf3, and Ova, does not impact germ cell behavior.** Adult testes reared at 25°C immunostained with anti-Vasa (red) and DAPI (blue) to display germ cells or nuclei respectively. Vasa and DAPI shown alone in black and white (A’-D’ and A’’-D’’, respectively). The GSC niche at the testis apex is marked with an asterisk. Scale bar = 70 μm. **(A)** 7-10 day c587-Gal4/+ control testes have undifferentiated germ cells with spherical morphology and strong DAPI staining near the niche, and weaker DAPI stain further down the testis coil. 0-3 day **(B)** c587-Gal4; UAS-CG4318-RNAi, **(C)** c587-Gal4; UAS-Taf3-RNAi, and (D) c587-Gal4; UAS-Ova-RNAi testes are similar to control. n > 20 for all genotypes.

