## Supplementary material for "Chigno/CG11180 and SUMO are Chinmo-Interacting Proteins with a Role in *Drosophila* Testes Stem Cells": Strain Table 1

| S1. Strains and plasmids used in this study. | | |
|---|---|---|
| **Name** | **Pertinent Genotypes or Parent Strains** | **Plasmids** |
| **YOK 1221** | AH109 (Clontech Laboratories Inc.) MATa, trp1-901, leu2-3, 112, ura3-52, his3-200, gal4Δ, gal80Δ, LYS2 : : GAL1UAS-GAL1TATA-HIS3, GAL2UAS-GAL2TATA-ADE2, URA3 : : MEL1UAS-MEL1 TATA-lacZ | BD-PGBKT7 -p53/(matα); AD-Y187-pGADT7-T (matα) |
| **YOK 3361** | AH109 | BD-Chinmo Native; AD-SMT3 |
| **YOK 3362** | AH109 | BD-Chinmo Stop; AD-SMT3 |
| **YOK 3392** | AH109 | BD-Chinmo Native; AD-Chinmo Native |
| **YOK 3394** | AH109 | BD-Chinmo Native; AD-CG5694 |
| **YOK 3396** | AH109 | BD-Chinmo Native; AD-CG4318 |
| **YOK 3397** | AH109 | BD-Chinmo Native; AD-CG2009/TAF3 |
| **YOK 3399** | AH109 | BD-Chinmo Native; AD-CG18269 |
| **YOK 3401** | AH109 | BD-Chinmo Native; AD-CG11180/Chigno |
| **YOK 3406** | AH109 | BD-Chinmo Native; AD-Empty |
| **YOK 3417** | AH109 | BD-Chinmo Non-Stop; AD-SMT3 |
| **YOK 3428** | AH109 | BD-Empty; AD-SMT3 |
| **YOK 3429** | AH109 | BD-Slx5; AD-SMT3 |
| **YOK 3517** | AH109 | BD-Chinmo Native; AD-Chinmo Stop |
| **YOK 3518** | AH109 | BD-Chinmo Non-Stop; AD-Chinmo Stop |
| **YOK 3519** | AH109 | BD-Chinmo Stop; AD-Chinmo Stop |
| **YOK 3520** | AH109 | BD-Chinmo Native; AD-Chinmo Native |
| **YOK 3521** | AH109 | BD-Chinmo Non-Stop; AD-Chinmo Native |
| **YOK 3522** | AH109 | BD-Chinmo Stop; AD-Chinmo Native |
| **YOK 3795** | AH109 | BD-SMT3; AD-Chigno WT |
| **YOK 3797** | AH109 | BD-SMT3; AD-Chigno sim* |
| **YOK 3799** | AH109 | BD-Chinmo Native; AD-Chigno WT |
| **YOK 3801** | AH109 | BD-Chinmo Native; AD-Chigno sim* |
| **YOK 3803** | AH109 | BD-Empty; AD-Chigno WT |
| **YOK 3805** | AH109 | BD-Empty; AD-Chigno sim* |
| **YOK 3813** | AH109 | BD-Empty; AD-Chinmo Stop |

**Table S1. Strains and plasmids used in this study.**

| Name | Plasmids |
|---|---|
| **BOK 293** | BD-Slx5 |
| **BOK 295** | BD-SMT3 |
| **BOK 312** | pOAD vector only (AD) |
| **BOK 313** | pOBD vector only (BD) |
| **BOK 571** | AD-SMT3 |
| **BOK 1311** | BD-Chinmo Native |
| **BOK 1315** | AD-Chinmo Native |
| **BOK 1316** | BD-Chinmo Stop |
| **BOK 1317** | AD-Chinmo Stop |
| **BOK 1339** | AD-CG4318 |
| **BOK 1340** | AD-CG18269 |
| **BOK 1341** | AD-CG11180/Chigno WT |
| **BOK 1342** | AD-CG11180/Chigno |
| **BOK 1350** | AD-CG2009 |
| **BOK 1353** | AD-CG5694 |
| **BOK 1376** | BD-Chinmo Non-Stop |
| **BOK 1626** | AD-Chigno sim* |
