## Supplementary figures and images for "Chigno/CG11180 and SUMO are Chinmo-Interacting Proteins with a Role in *Drosophila* Testes Stem Cells"

### Fig.S1 Rinehart

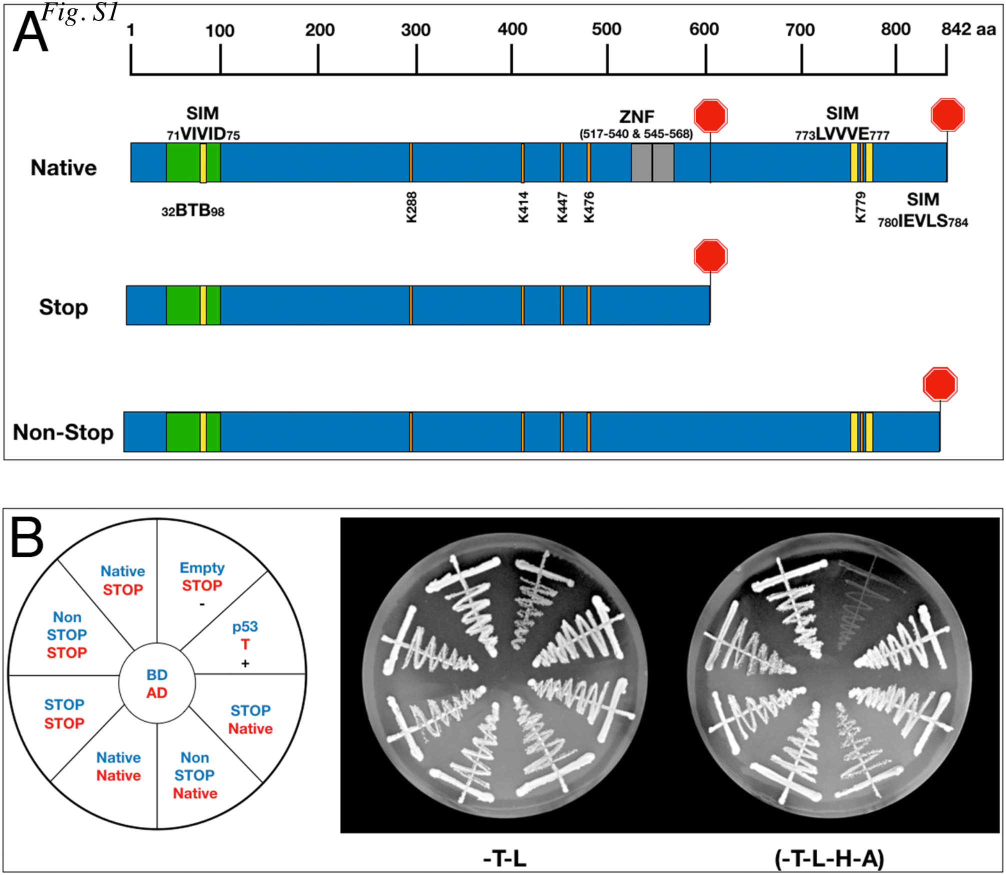

### Fig.S2 Rinehart

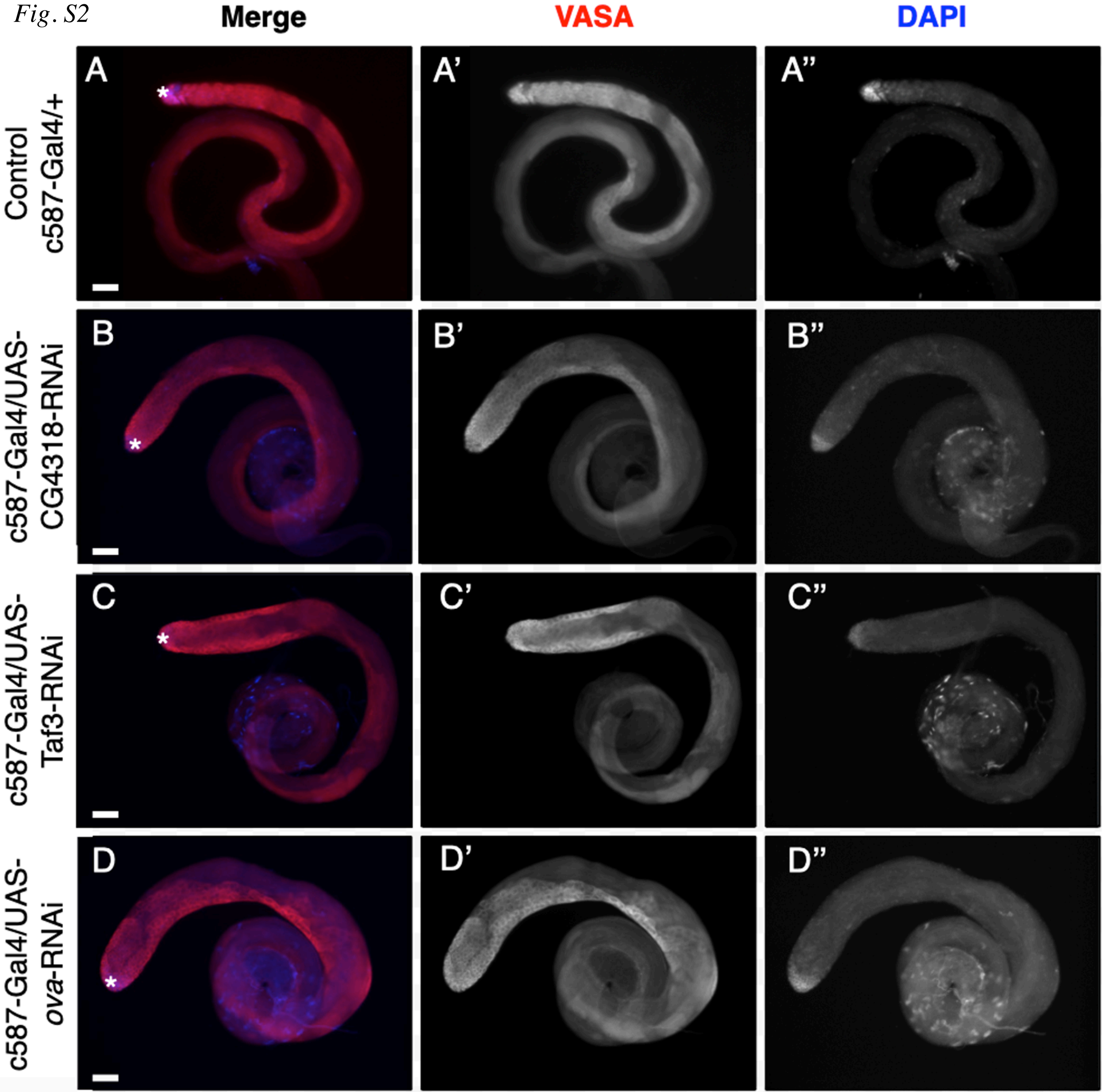
